# Host genetics and COVID-19 severity: increasing the accuracy of latest severity scores by Boolean quantum features

**DOI:** 10.1101/2023.02.06.527291

**Authors:** Gabriele Martelloni, Alessio Turchi, Chiara Fallerini, Andrea Degl’Innocenti, Margherita Baldassarri, GEN-COVID Multicenter study, Simona Olmi, Simone Furini, Alessandra Renieri

## Abstract

The impact of common and rare variants in COVID-19 host genetics is widely studied in [16]. Here, common and rare variants were used to define an interpretable machine learning model for predicting COVID-19 severity. Firstly, variants were converted into sets of Boolean features, depending on the absence or the presence of variants in each gene. An ensemble of LASSO logistic regression models was used to identify the most informative Boolean features with respect to the genetic bases of severity. After that, the Boolean features, selected by these logistic models, were combined into an Integrated PolyGenic Score, the so called IPGS, which offers a very simple description of the contribution of host genetics in COVID-19 severity. IPGS leads to an accuracy of 55-60% on different cohorts and, after a logistic regression with in input both IPGS and the age, it leads to an accuracy of 75%. The goal of this paper is to improve the previous results, using the information on the host organs involved in the disease. We generalized the IPGS adding a statistical weight for each organ, through the transformation of Boolean features into “Boolean quantum features”, inspired by the Quantum Mechanics. The organs’ coefficients were set via the application of the genetic algorithm Pygad and, after that, we defined two new Integrated PolyGenic Score (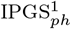 and 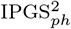). By applying a logistic regression with both 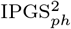 (or indifferently 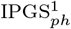) and age as input, we reach an accuracy of 84-86%, thus improving the results previously shown in [16] by a factor of 10%.

## 1. Introduction

In early December 2019 the Coronavirus appeared in Wuhan, China. On January 9th, 2020, the Chinese Center for Disease Prevention and Control (CDC) reported that a new Coronavirus (initially called 2019-nCoV and now called SARS-CoV-2) has been identified as the causative agent and its genomic sequence has been published. The appearance of new pathogenic viruses for humans, previously circulating only in the animal world, is a widely known phenomenon, called spill over, and it is believed that it is at the origin of the SARS-CoV-2. Despite the drastic, large-scale containment measures promptly implemented by the Chinese government, in a matter of a few weeks the disease had spread well out-side China, reaching countries in all parts of the globe. The Oms Director-General Tedros Adhanom Ghebreyesus announced the name for the disease on February 11, 2020, during the extraordinary press conference dedicated to the virus: the disease has been named as “COVID-19”, where “CO” stands for corona, “VI” for virus, “D” for disease and “19” indicates the year in which it firstly occurred. Finally, on March 11th, 2020 Oms declared that COVID-19 can be defined as a pandemic.

COVID-19 disease, due to its rapid worldwide spreading, has lead to the most severe pandemic since the deadly Spanish flu, which killed up to 100 million individuals in the past century. Most COVID-19 affected patients have mild symptoms, but around 20% of cases need hospitalization, ranging from severe to critical illness requiring very intensive help. Patients with severe illness are often older and/or have comorbidities (e.g. cardiovascular or chronic respiratory disease, diabetes, hypertension and cancer). Moreover the organ involvement turned out to be related to disease severity, even though the correlation is still under clarification [12, 4], while another factor that ended up being discriminant is sex, with males tending to have a more severe disease respect to females [37]. However, these factors do not fully explain the differences in severity.

It is now well recognized that host genetic factors play a fundamental rule in COVID-19 clinical outcome. Different pathogenetic mechanisms can be involved as new genetic predisposing factors emerge, such as different immunogenicity/cytokine production capability, as well as receptor permissiveness to virus and antiviral defenses. However, with very few genetic factors identified up to now, we are still very far from understanding the real relevance of host genetics. The better understanding of host genetic factors is fundamental to predict patients who are at risk of severe disease and prevent and/or offer personalized and efficient treatments. Moreover, novel genetic discoveries could also inform therapeutic targets for drug repurposing. A pivotal example has been the discovery of homozygous deletions in the CCR5 gene conferring resistance to HIV-1 infection, which led to the development of a drug that successfully made it through clinical trials [21].

Traditional methods for assessing the genetic bases of complex disorders include Genome-Wide Association Studies (GWASs) for common variants and Burden test for rare variants. GWASs focus mainly on common variants and are based on a straightforward comparison frequency of about 700,000 genomic single-nucleotide polymorphisms (SNPs) in cases/controls (mostly noncoding). The coverage of the coding SNPs is usually performed throughout imputed data, e.g., imputing 2 million SNPs from 700k SNPs by Linkage Disequilibrium. The method is based on multiple independent tests and has a high threshold for significance. Moreover GWASs require sample sizes of ten-hundred thousand subjects [33, 20, 23, 32]. On the other hand the Burden test is based on an aggregation of rare, protein-altering variants and a comparison between cases and controls. The reasoning behind the Burden test is that grouping variants with a large effect size at a gene level might improve power. Like GWASs, the Burden test method needs hundreds of thousands of participants to detect statistically significant associations [22]. These methods have been employed for many years but failed to fully unravel the complexity of human traits. Complex disorders such as COVID-19 are expected to be regulated by thousands of genes with different weights of contribution [28, 6]. Indeed, in common genetic diseases such as cardiovascular or neurodegenerative disorders, the identified genetic markers were not sufficient for full use in clinical practice to predict and treat the disease.

To overcome these limitations, the interplay between host genetics, computational statistics and dynamic system theory is necessary. Even though the scientific community has made a big effort to analyze the epidemic data made available by the Center for Systems Science and Engineering at Johns Hopkins University [13], the applications of mean-field models able to predict the kinetics of the epidemic spreading [29, 30, 25, 9, 7, 18, 17, 1, 10, 26] cannot help identifying the gene variants that determine the risk of severity in order to understand the pathophysiological mechanisms responsible for severe disease in heterogeneous groups of patients. At the contrary, Machine Learning (ML) approaches offer an innovative tool for managing complex problems by significantly increasing our capacity to identify complex patterns of variations. Using data from the Whole Exome Sequencing (WES), a first line of ML method, i.e. a LASSO logistic regression, has been applied to extract some thousands of coding genetic features contributing to COVID-19 severity [35, 16]. Subsequent functional validation of extracted features demonstrated that, in each tested case, the association with severity has a biological basis and suggested hints for adjuvant treatment [5, 15, 14, 11, 3, 2, 27, 31]. Using the extracted features it has been built a severity score named Integrated PolyGenic Score (IPGS), whose performances reached about 75% for both sensitivity and specificity [16]. In this contribution we want to improve the severity score performances, with the aim of increasing both metrics and the understanding of bio-molecular mechanisms for personalized treatment using innovative ML methods. In the following we present two severity scores that takes into account the phenotype of the analyzed patients, i.e. the set of their observable characteristics or traits. In particular we take into account the involvement of single organs in the development of the COVID-19 disease and the age of patients when they contract the virus. More specifically, the Methods section is devoted to the description of the implemented severity scores and the applied methods. In Sec. 3 are presented the performances of the new severity scores with respect to the IPGS, while a discussion on the presented results is reported in section Discussion.

## 2. Methods

### 2.1. Data collection

Clinical data have been collected through a Clinical questionnaire containing more than 160 clinical items. Items concerning organ/system involvement (heart, liver, pancreas, kidney, and olfactory/gustatory and lymphoid systems) have been synthesized in a binary mode, where 1 means standard medical parameters indicating specific organ involvement (respiratory severity, taste/smell involvement, heart involvement, liver involvement, pancreas involvement, kidney involvement, lymphoid involvement, blood clotting, cytokines trigger and number of comorbidities like asthma, cancer, diabetes, dyslipidemia, hypertension, hypothyroidism or obesity) and 0 absence of involment of a certain organ/system. In this way clinical data are easily accessible for a rapid statistical analysis [12].

#### Phenotype definitions

The COVID-19 severity has been assessed using a modified version of the WHO COVID-19 Outcome Scale (COVID-19 Therapeutic Trial Synopsis 2020); specifically six classification levels have been used to code for the severity: (5) death; (4) hospitalized receiving invasive mechanical ventilation; (3) hospitalized, receiving continuous positive airway pressure or bilevel positive airway pressure ventilation; (2) hospitalized, receiving low-flow supplemental oxygen; (1) hospitalized, not receiving supplemental oxygen; and (0) not hospitalized. Through the application of the presented severity scores, this six-level classification (termed GRADING_5_) will be reduced to two different classifications: i) a binary classification of patients into mild and severe cases (termed GRADING_2_), where a patient is considered severe if hospitalized and receiving any form of respiratory support (WHO severity-grading equal to 4 or higher in 6 points classification); ii) a 3-level classification (termed GRADING_3_), where the patients are classified into non-hospitalized (WHO severity-grading equal to 0, 1), hospitalized not receiving supplemental oxygen or receiving low-flow oxygen (WHO severity-grading equal to 2, 3) and patients with severe disease (WHO severity-grading equal to 4 or higher).

#### GEN-COVID cohort

About 2,000 patients have been sequenced by Whole-Exome Sequencing (WES) within the GEN-COVID Multicenter Study and already included in the model described in [16]. WES with at least 97% coverage at 20x was performed using the Illumina NovaSeq6000 System (Illumina, San Diego, CA, USA). Library preparation was performed using the Illumina Exome Panel (Illumina) according to the manufacturer’s protocol. Library enrichment was tested by qPCR, and the size distribution and concentration were determined using Agilent Bioanalyzer 2100 (Agilent Technologies, Santa Clara, CA, USA). The Novaseq6000 System (Illumina) was used for DNA sequencing through 150 bp paired-end reads. Variant calling was performed according to the GATK4 (O’Connor and Auwera 2020) best practice guidelines, using BWA (Li and Durbin 2010) for mapping and ANNOVAR (Wang et al. 2010) for annotating.

### 2.2. Post-Mendelian paradigm for COVID-19 modelization

In [16] some of the present authors have developed an easily interpretable model that could be used to predict the severity of COVID-19 from host genetic data. Patients were considered severe when hospitalized and receiving any form of respiratory support. The focus on this target variable is motivated by the practical importance of rapidly identifying which patients are more likely to require oxygen support, in an effort to prevent further complications. The complexity of COVID-19 immediately suggests that both common and rare variants are expected to contribute to the likelihood of developing a severe form of the disease. However, the weight of contribution of common and rare variants to the severe phenotype is not expected to be the same. A single rare variant that impairs the protein function might cause a severe phenotype by itself after viral infection, while this is not so probable for a common polymorphism, which is likely to have a less marked effect on protein functionality. These observations led to the definition of a score, named Integrated Polygenic Risk Score (IPGS), that includes data regarding the variants at different frequencies:

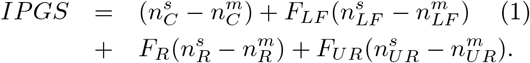

In this equation, the *n* variables indicate the number of input features of the predictive model that promote the severe outcome (superscript s) or that protect from a severe outcome (superscript m) and with genetic variants having Minor Allele Frequency (MAF)≥ 5% (common, subscript C), 1%<MAF5 ≤% (low-frequency, subscript LF), 0.1%<MAF≤1% (rare, subscript R), and MAF≤0.1% (ultra-rare, subscript UR). The features promoting or preventing severity were identified by an ensemble of logistic models. The weighting factors *F_LF_*, *F_R_*, and *F_UR_* model the different penetrant effects of low-frequency, rare, and ultra-rare variants, compared to common variants (for which the weighting factor has been chosen equal to 1). Thus, the four terms of Eq. (1) can be interpreted as the contributions of common, low-frequency, rare, and ultra-rare variants to a score that represents the genetic propensity of a patient to develop a severe form of COVID-19. In particular, note the difference in sign between the severe and mild variants, which respectively represent a predisposing factor compared to a protection factor. The model including IPGS exhibited an overall accuracy of 73%, precision equal to 78%, with a sensitivity and specificity of 72% and 75%, respectively, thus showing a statistically significant increase in the performances with respect to logistic models that adopt as input features only age and sex. However, in order to design prevention and treatment protocols in view of personalized medicines development, the predictability of the post-Mendelian paradigm for COVID-19 modelization should be further increased.

### 2.3. First phenotype-based IPGS 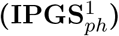

To improve the ability of the IPGS to predict the severity of the disease, while keeping the linearity of the formula, we first pass to a vectorial formulation, where both the Boolean variables of the individual patients and the Boolean variables of the single variants are tranformed into vectors with components 0, 1. To each patient and to each single variant is associated a vector, which has univocally defined non-zero component: the non-zero components of the patient vector *P_i_* and the variants vector 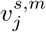, allow us to codify the situation of each patient, that has a unique set of variants and a specific clinical condition when he/she has contracted the COVID-19 disease. Specifically, the clinical overview takes into account the involment of the organs for each subject, that are included in the matrix **O**, whose entries *O_ij_* are 1 (0) in case the organ *j* is involved (non involved) in the disease development of patient *i*. A scalar product between the vector of the single patient and the vectors of the genetic variants through the matrix of the organs univocally identifies the phenotypic characteristics of the patients, weighted by the variants. Finally, we release the condition that mild variants always protect from a severe outcome, thus being subtracted in the Eq. (1) and we do not fix a priori the sign of the mild variants. Starting from a vectorial formulation of the severity score, we are now able to write down a severity score that includes non only the genetic features of the single patients, but also the involvement of the organs in the disease development, through the matrix of the organs *O_ij_*. The score index that encompasses the phenotypical characteristics of the patients is called 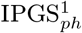 and it reads as

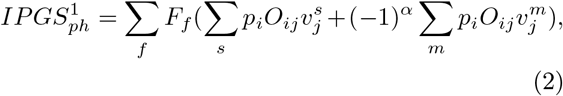

where *F_f_* is the coefficient representing the frequency of the variants, as in Eq. (1), and the subscript *f* identifies either common, low frequency, rare and ultrarare variants. As introduced before, *p_i_* represents the single patient vector, while 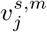 represents the vector of severe or mild variants, where we can distinguish between severe and mild according to the superscript. Differently from Eq. (1), we leave not fixed the sign of the variants, therefore in the sum over the mild variants, the sign remains a coefficient to be fitted through the parameter *α*. This results in having 17 more parameters to be fixed. Some examples of Eq. (2) are reported in Sec. 1 in the Supplementary Information; specifically are reported some case examples for different involved organs and different genetic features.

### 2.4. Second phenotype-based IPGS 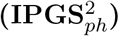

Inspired by quantum mechanics we try to generalize here the severity score presented in Eq. (2), esplicity introducing in the formula the age of each patient and leaving the possibility, thanks to the quantum mechanics formalism, to introduce into the new severity score expression, more general phenotype definitions. For a brief introduction to the quantum mechanics formalism see Sec. 2 of the Supplementary Information. Borrowing the formalism of quantum mechanics, we use the following elements to construct the second severity score 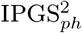:

- the patient is described in terms of a vector |*p* >, which represents a state in quantum mechanics and describes the condition of the single human being;
- the genetic variants are also expressed in terms of vectors 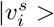, which represent a vector’s basis to calculate the expectation value of the physical observables;
- the organs can be considered as the physical observable O, whose expectation value represents our quantum-like 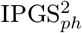;
- the time, related to the evolution operator, represents the patient’s age;
- the mild or severe variants can be represented through a spin variable *s* which takes values 1/2 or −1/2.

In order to better clarify the role played by each single element in the severity score, we explicity write down the values we assign to the new Boolean variables. More in detail, we can distinguish the state of the single *i* – *th* patient via assigning a sequence of values 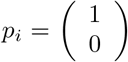 or 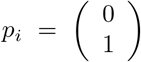. Since we are dealing with patients that have contracted COVID19, but have different phenotypic characteristics (i.e. different organs involved in the disease course), the sequence of 2-dim vectors with 0, 1 values is unique for each patient and it allows us to select the right organ involment when performing a scalar product. To gain a better insight on the construction of the severity score we refer to Sec. 1 in the Supplementary Information. Similarly, the same concept is reported on the genetic variants: if the patient shows the *j* – th variant, the vector 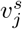 takes the values 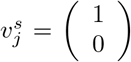, otherwise we assign 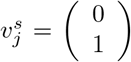. We thus have constructed the quantum-like Boolean variables (or features) and we are ready to define the mathematical structure of the 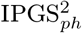:

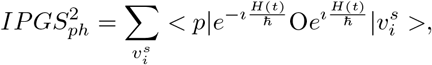

where *H*(*t*) is the Hamiltonian operator and 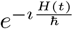 represents the time-evolution operator. To make the previous formula manageable, we perform some approximations, first of all inserting a completeness of the vectors of our base 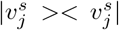, which represents the genetic heritage of the human being:

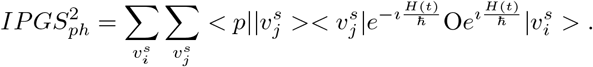

We can perform the subsequent approximations along two different lines: either i) we suppose that the vectors of the variants are eigenvalues of the Hamiltonian *H*(*t*), or ii) we perform the infinite time limit of the system. In the first case, if we assume that *E_i_* represents the eigenvalue of the hamiltonian *H*(*t*) related to the precise state 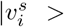, we can approximate 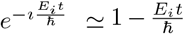. *E*, corrisponding in general to the total energy of the system, can be put in correlation with the comorbidity of the system human being. In this case we obtain

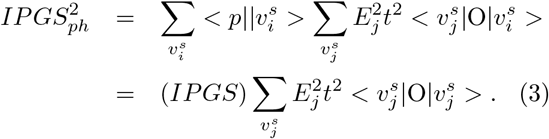

In the latter case, the limit *t* → ∞ corresponds to the assumption that the patient has contracted the COVID19 and his/her status is characterized by a small number of variants that are only those relevant to the contraction/development of the disease. The small set of variants that are related to the disease and influence the clinic outcome of the patients can be called *variants of the saddle point* [8] and identified with 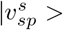. In this case the severity score reads as

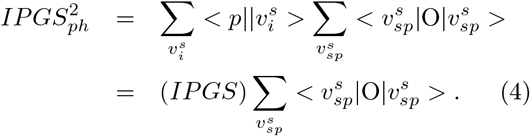

In both Eqs. (3), (4), it is present the term 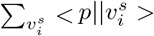, that represents the scalar product between the vector that identifies the patients’ clinical state and the vector taking into account the genetic variants. Thanks to the characterization of the single genetic variant in terms of the spin variable *s* (*s* = mild, severe), this scalar product constitutes the IPGS previously defined in Eq. (1). In other words, the scalar product 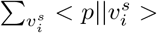 is the overlap between the initial state, i.e. the state of the patient and the base of our system (the host genetics).

Here the severity score Eq. (1) turns out to be corrected by a form factor that constitutes either the expectation value of the organs on the state of all genes, weighted with the age in Eq. (3), or the interplay between the variants of the genes, known to be associated to viral susceptibility and disease severity, and the patient status in Eq. (4). While the form factor present in Eq. (3) can be easily interpreted as the clinical status of the patient, where organs correlate with the genetic variants, the form factor in Eq. (4) has a more complex interpretation. Somehow the vector 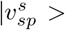 represents the variants selected by the LASSO regression in [16] and Eq. (4) can be interpreted as the product between the scores previously defined in Eqs. (1), (2): 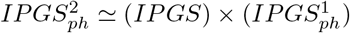.

Irrespectively of the fact that we have just taken inspiration from quantum mechanics, since in the previous definitions of 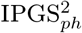 there is no real time evolution and the vectors are fixed a priori, as well as the structure of the observables, using the quantum mechanics formalism helped us to generalize the problem and build a severity score that, in principle, can be generalized to other diseases.

### 2.5. The genetic algorithm Pygad

The genetic algorithm is a method for solving both constrained and unconstrained optimization problems that is inspired by natural selection, the process that drives biological evolution. In a genetic algorithm, we start with an initial population of chromosomes, which are possible solutions to a given problem. Those chromosomes consists of an array of genes whose values vary in a pre-defined range. The whole optimisation problem is encoded into a fitness function, which receives a chromosome and returns a number that tells the fitness (or goodness) of the solution. The higher the fitness, the better the solution encoded in the chromosome. The genetic algorithm repeatedly modifies a population of individual solutions. At each step, the genetic algorithm selects individuals from the current population to be parents and uses them to produce the children for the next generation. At each iteration (generation), a number of good chromosomes are selected for breeding (parent selection). Parents are combined two-by-two (crossover) to generate new chromosomes (children). The children are finally mutated by (randomly) modifying part of their genes, allowing for completely new solutions to emerge. Over successive generations, the population “evolves” toward an optimal solution, as it is shown in the flow chart of a Genetic Algorithm (GA) in Fig. 1.

**Figure 1:**
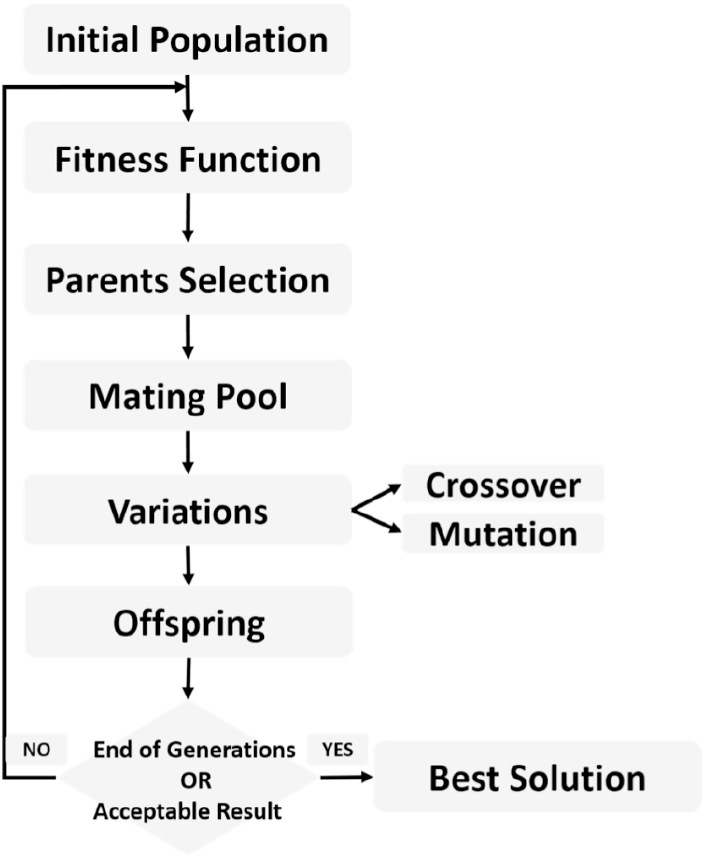
The Flow chart of a Genetic Algorithm

The genetic algorithm is usually applied to solve problems in which the objective function is discontinuous, nondifferentiable, stochastic, or highly nonlinear. Among the genetic algorithms, we find PyGAD, an open-source Python library [19], which supports a wide range of parameters to give the user control over everything in its cycle of operations (see Fig. 2).

**Figure 2:**
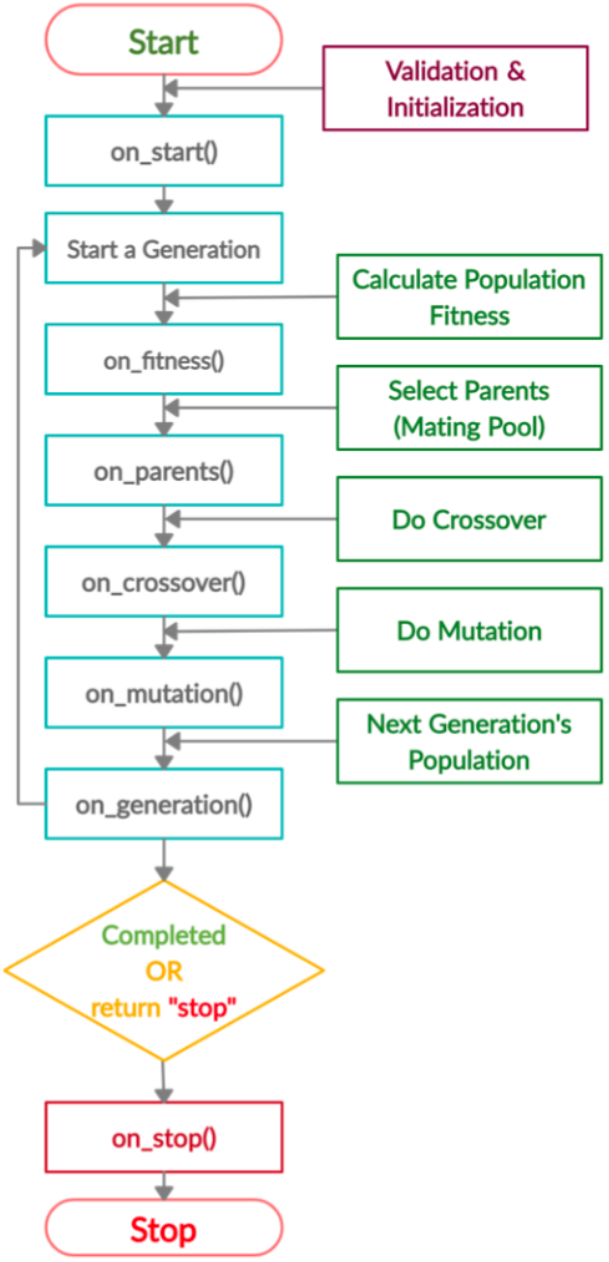
PyGAD lifecycle

#### Testing and training procedure

The dataset was randomly divided into a training set and a test set (50/50) for a total of 2240 patients. The PyGAD algorithm has been implemented with the following characteristics in order to converge to a stable solution:

- Number of solutions (i.e. chromosomes) within the population=32
- Number of generations 250-500
- Number of solutions to be selected as parents= 8
- Parent selection type = sss (for steady-state selection). In sss case only a few individuals are replaced at a time, meaning most of the individuals will carry out to the next generation.
- Number of parents to keep in the current population = 1
- Crossover operation = single_point (for singlepoint crossover). All genes to the right of that point are swapped between the two parent chromosomes. This results in two offsprings, each carrying some genetic information from both parents.
- Type of the mutation operation = random (for random mutation)
- The probability of selecting a gene for applying the mutation operation = 0.2 (For each gene in a solution, a random value with probability 20% is generated)

In most part of the developed training/testing tests, the number of generations able to garantee a convergence of the solution is 250. We considered as a converged solution, the one that has reached an asymptotic value within the duration of the test.

The training/testing procedure, for each severity score, has been implemented separately on the male and female datasets. The whole procedure is made up of two parts, both used on the testing and training samples. In the first part we let the genetic algorithm run over the training sample to fit the parameters of the severity scores Eqs. (2), (3), that produces the best estimate of the *N*-level classification of the patient severity (i.e. GRADING_N_ parameter). Then the 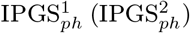 is computed over the test sample, by employing the fitted parameters. For each severity score and each dataset, the training and testing tests has been repetead 10 times. Since the mutation process is random, this is done to ensure that we are able to get the bes solution among a sufficient number of iterations. In the second part of the procedure a multivariable logistic regression is fitted using the 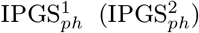 computed according to the steps described above, together with other input parameters (age, IPGS and sex), to predict the same GRADING_N_ parameter. The logistic model is first trained on the training sample and then tested on the test sample. The solutions that are shown in the following section, are those corresponding to the best performances among the obtained results.

## 3. Results

The severity scores Eqs. (2), (3) are used, together with the GRADING_N_ data, for training a model that predicts the COVID-19 severity. In particular, the training procedure is devoted to fitting the parameters that are present in the severity score equations: 17 free parameters for Eq. (2) and 18 free parameters for (3) respectively. Fitting the parameters will allow us to assess, for each patient, the level of severity of his/her COVID-19 infection, in terms of 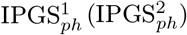. Since the final goal is to produce a N-level classification of the patient severity, we have to further reduce the results obtainable from Eqs. (2), (3), in a N-level classification along the line of GRADING_N_.

To obtain the best possible fit we have implemented the genetic algorithm Pygad with the following step fitness function:

- We assign a reward 50 in case the obtained score value is 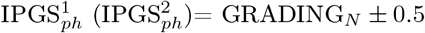
- We assign a reward 5 in case the obtained score value is 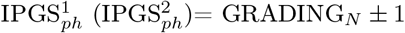
- We assign 0 otherwise.

First we present the results related to GRADING_2_, where we have reduced the severity scores to a binary classification of patients into mild and severe cases, considering a patient severe (GRADING_2_=1) if hospitalized and receiving any form of respiratory support or healthy (GRADING_2_=0) in all the other cases. Furthermore, a multivariable logistic regression was fitted using as possible inputs 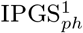 and 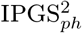, alone or combined with IPGS, age and sex. In Fig. 3 are shown the confusion matrices, also known as error matrices [36], for the male (panels (a), (b)) and female (panels (c),(d)) data set, where the best fit is presented for both sets. The performances of the logistic regression increase when multiple predictor variables are used, instead of the single severity score 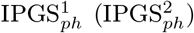. In particular the best fit is obtained, both for the male and female data set, when using in input age, IPGS and 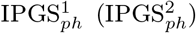. While for males the new severity score 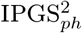 gives more accuracy than 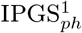, for females the logistic regression gives higher accuracy when giving in input age, IPGS and 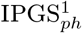, with respect to age, IPGS and 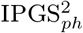. However the reached accuracy values are comparable for both severity scores; specifically we reach an accuracy> 86% for the male data set, and an accuracy > 83% for the female data set. Moreover the confusion matrices indicate that we are able to predict the grading 1 with a reasonably high successfully rate, while we have more difficulties in predicting the grading 0. Most errors are done, both for the female and male data sets, when the actual score is 0, but we predict 1.

**Figure 3:**
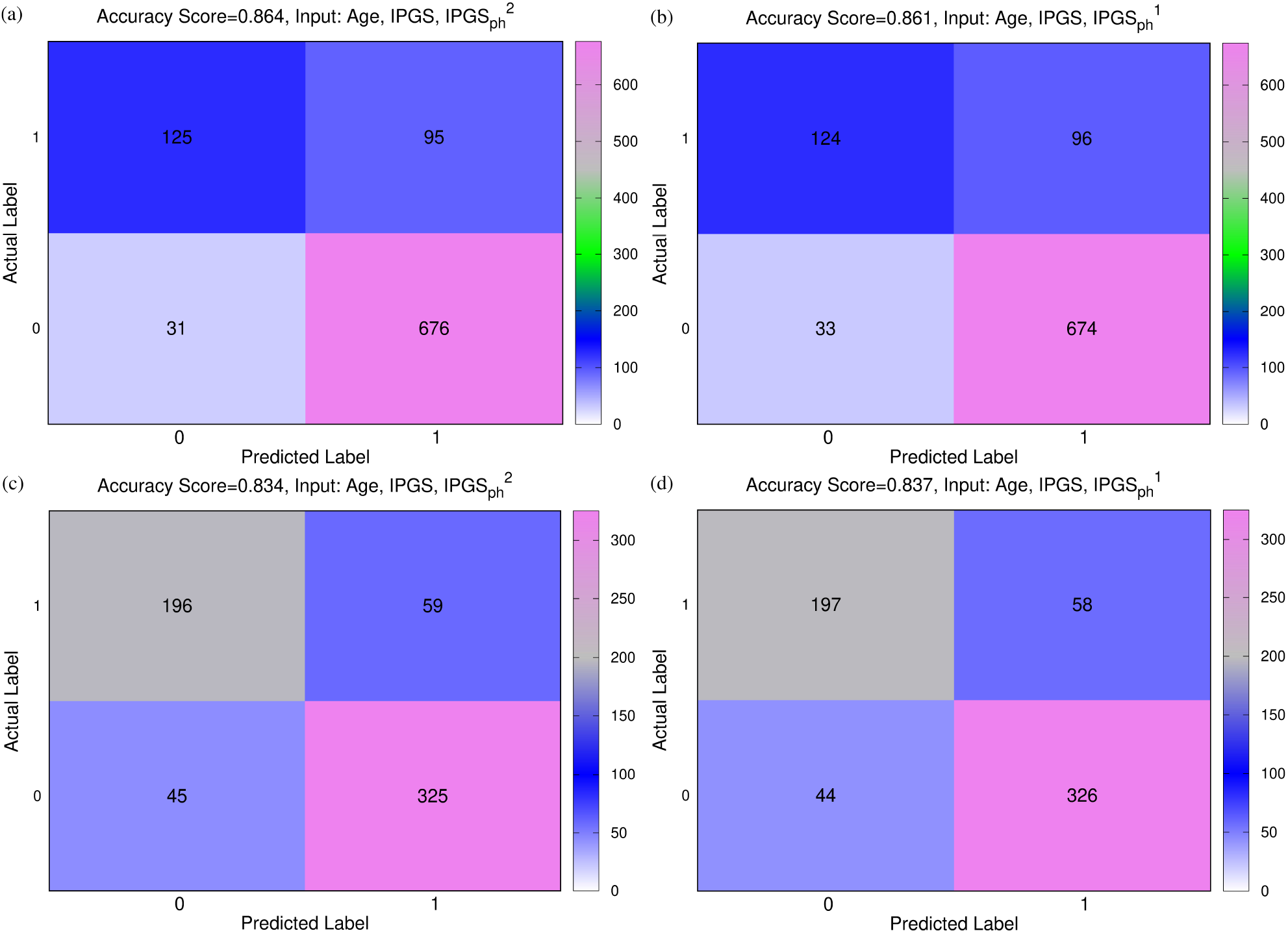
(a), (b) Confusion Male Matrix from Logistic Regression. (a) Input: Age, IPGS, 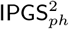. Accuracy=86,4%. (b) Input: Age, IPGS, 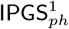. Accuracy=86,1%. (c), (d) Confusion Female Matrix from Logistic Regression. (c) Input: Age, IPGS, 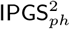. Accuracy=83,4%. (d) Input: Age, IPGS, 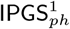. Accuracy=83,8%.

In order to confirm the godness of the results previously shown, we evaluate the increase of the performances of the severity score Eqs. (2), (3), with respect to the performances of a model where the values of the 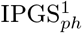 feature have been shuffled. In other words, we recalculate the 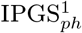 by assigning to each patient a random distribution of variants instead of his/her genetic variants. To compare the results we perform a logistic regression with in input age and 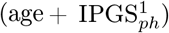 calculated with the shuffled variants (see Fig. 4 and Tables 1, 2). The performances of the logistic regression with in input age and 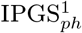 with shuffled variants are comparable with those obtained with only age in input, thus confirming that the calculation of the severity score with shuffled variants does not add any information with respect to the age. Moreover, in terms of accuracy, the score of the logistic regression shown in Fig. 4 both for males (panel a) and females (panel b), is lower than the corresponding one presented in Fig. 3. The accuracy for the 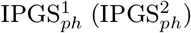 with the genetic variants is increased by a factor 10% (9%) for the male (female) sample with respect to the 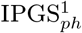 with shuffled variants. This means that the contribution of genetic variants to the information is fundamental in our analysis, in addition to the age factor that seems to be dominant in determining the severity of the disease.

**Figure 4:**
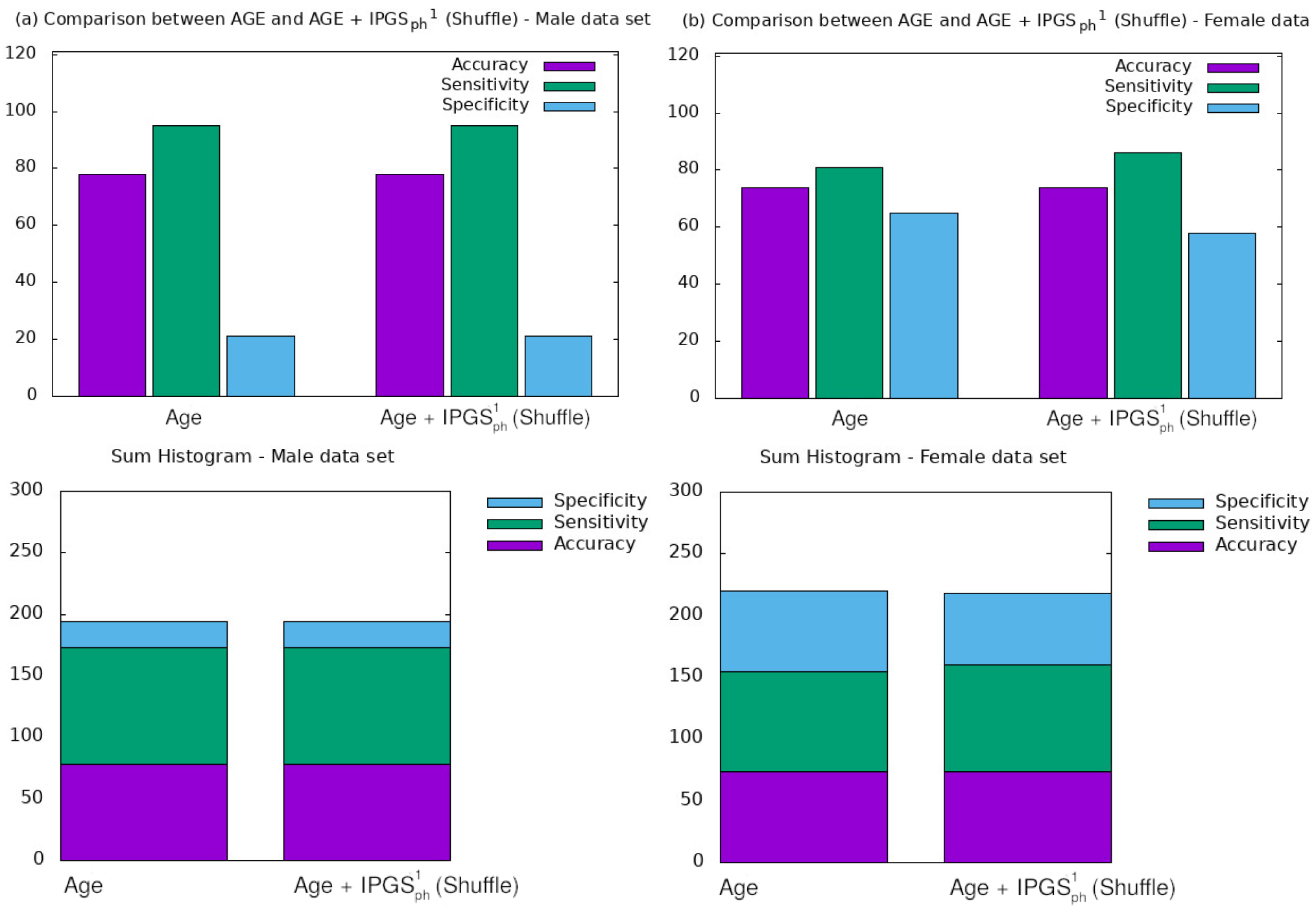
Comparison between the results obtained from a logistic regression with in input age or 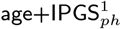 with shuffled variants for the (a) male sample and the (b) female sample.

**Table 1.** Accuracy, sensitivity and specificity scores resulting from a logistic regression on male data set for GRADING_2_.

| Input Variables | Accuracy | Sensitivity | Specificity |
| --- | --- | --- | --- |
| age | 0.78 | 0.95 | 0.21 |
| age+ IPGS<sub>ph</sub><sup>1</sup> (Shuffle) | 0.78 | 0.95 | 0.21 |

**Table 2.** Accuracy, sensitivity and specificity scores resulting from a logistic regression on female data set for GRADING_2_.

| Input Variables | Accuracy | Sensitivity | Specificity |
| --- | --- | --- | --- |
| age | 0.74 | 0.81 | 0.65 |
| age+ IPGS<sub>ph</sub><sup>1</sup> (Shuffle) | 0.74 | 0.86 | 0.58 |

In order to further investigate the role played by the age and other factors that seems to be discriminant, i.e. sex, in comparison with the new severity scores here presented, we report a comparison between the performances of the logistic regression when the predictor variables in input are (age + sex) or 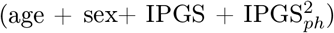 (see Fig. 5 and Table 3 for an overview). Here we report just the results for 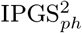, but comparable results are obtained when fitting the logistic regression with in input (age + sex) or 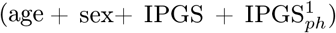. Considering both the role played by the genetic variants through the IPGS and the phenotypic information on the patients through the 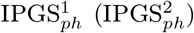, we observe an improvement of the sensitivity, specificity, accuracy scores with respect to the case where only the information on age and sex is used as input of the logistic regression. This confirms the initial hypotheses that comorbidities, age and sex are important to determine the disease severity, but these factors do not fully explain the differences in severity. More in detail, when comparing the results of the logistic regression with in input (age+sex+IPGS), 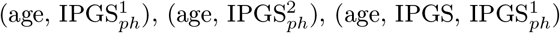 or 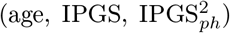 for the female (Fig. 6 (a)) and male (Fig. 6 (b)) data set, we observe that the best performances are obtained when using as input data age, IPGS and 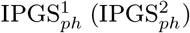. The numerical values correspondent to the histogram representation in Fig. 6 are reported in Tables 4, 5.

**Figure 5:**
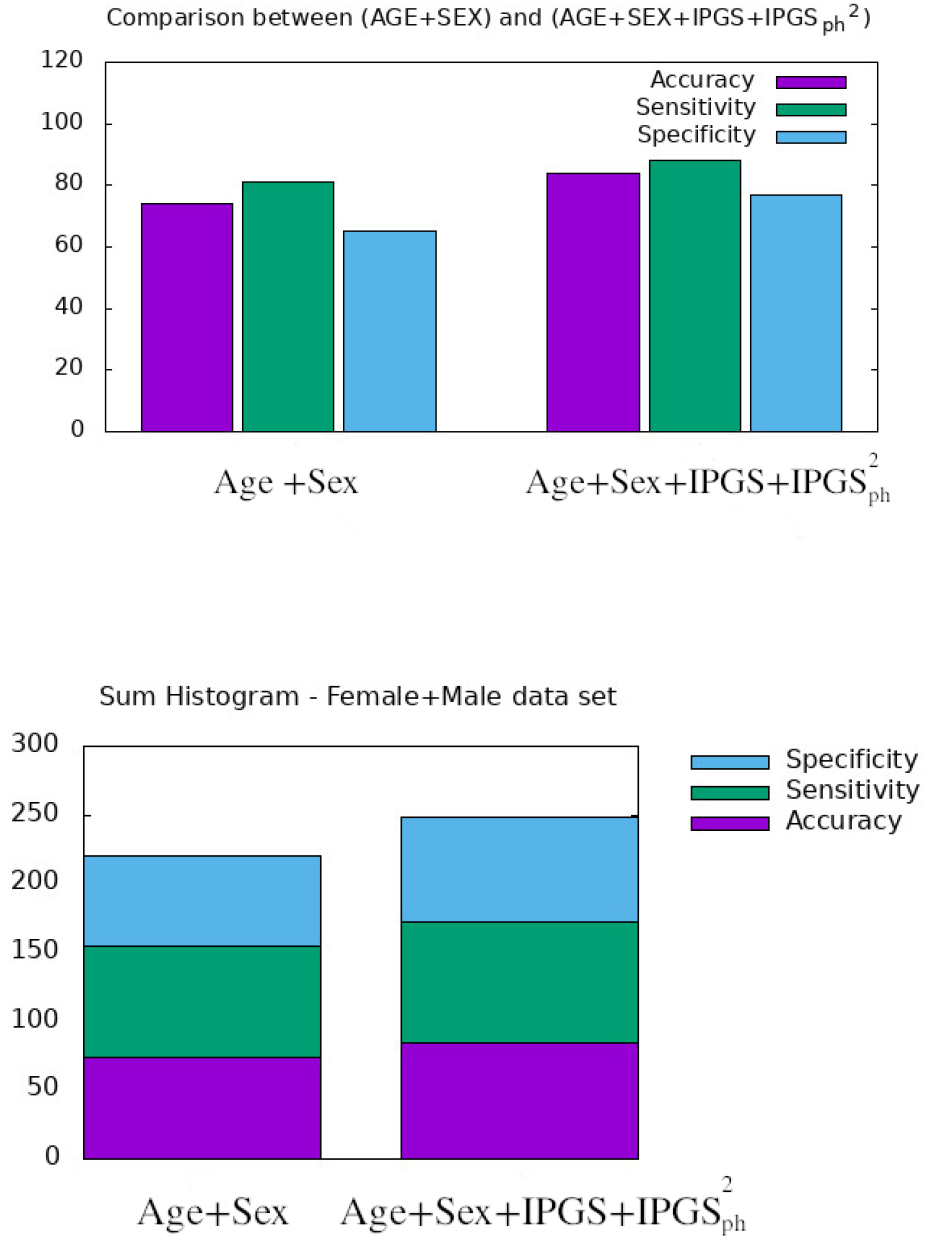
Comparison between AGE+SEX and 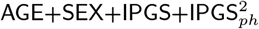 in input at the logistic regression on the total (female+male) data set.

**Figure 6:**
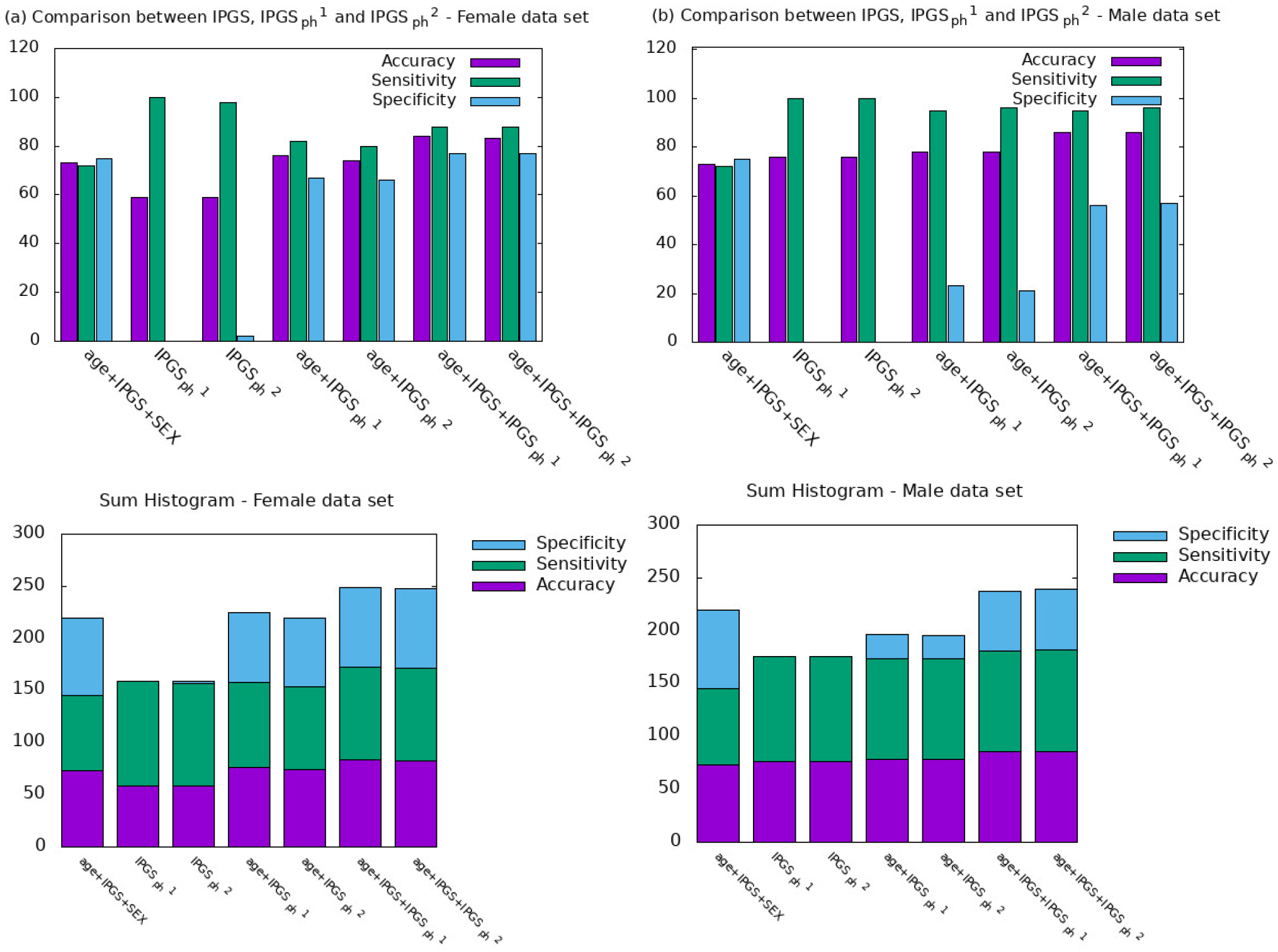
Comparison between IPGS, 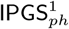 and 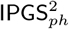 for the female (a) and male (b) sample.

**Table 3.** Accuracy, sensitivity and specificity scores resulting from a logistic regression on both female and male data set for GRADING_2_.

| Input Variables | Accuracy | Sensitivity | Specificity |
| --- | --- | --- | --- |
| age +SEX | 0.74 | 0.81 | 0.65 |
| age+ SEX+ IPGS + IPGS<sub>ph</sub><sup>2</sup> | 0.84 | 0.88 | 0.77 |

**Table 4.** Accuracy, sensitivity and specificity scores resulting from a logistic regression on the female data set for GRADING_2_.

| Input Variables | Accuracy | Sensitivity | Specificity |
| --- | --- | --- | --- |
| age + IPGS + SEX | 0.73 | 0.72 | 0.75 |
| IPGS<sub>ph</sub><sup>1</sup> | 0.59 | 1 | 0.0039 |
| IPGS<sub>ph</sub><sup>2</sup> | 0.59 | 0.98 | 0.016 |
| age+ IPGS<sub>ph</sub><sup>1</sup> | 0.76 | 0.82 | 0.67 |
| age+ IPGS<sub>ph</sub><sup>2</sup> | 0.74 | 0.80 | 0.66 |
| age+ IPGS + IPGS<sub>ph</sub><sup>1</sup> | 0.84 | 0.88 | 0.77 |
| age+ IPGS + IPGS<sub>ph</sub><sup>2</sup> | 0.83 | 0.88 | 0.77 |

**Table 5.**
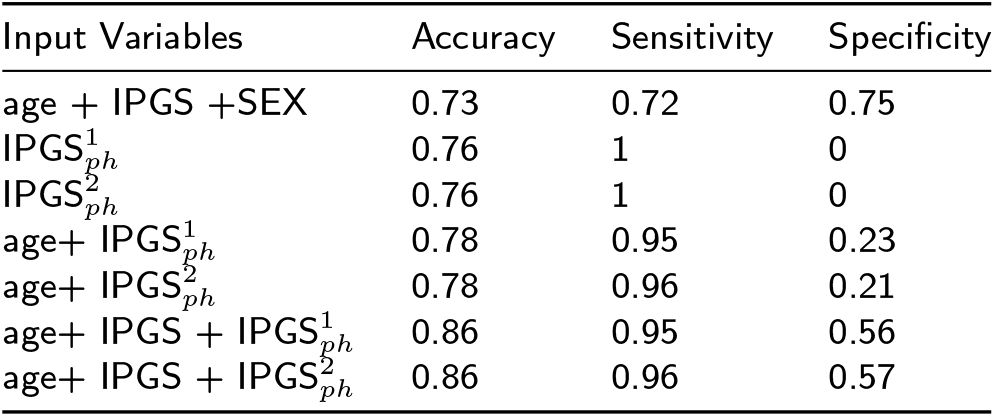
Accuracy, sensitivity and specificity scores resulting from a logistic regression on the male data set for GRADING_2_.

Therefore age, genetic variants and organ involvement seem to all concur and contribute to the amount of information necessary to reach good levels of sensitivity, specificity and accuracy scores (> 80% in all cases). Moreover, taking into account the organs involved during the disease, each with its statistical weight, leads to an improvement of the score of 10% compared to the previous work (corresponding to the case where age, sex and IPGS are used as input variables), in terms of forecast accuracy.

Finally we spend some words on the comparison between 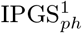 and 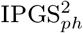 (see Fig. 6 and Tables 4, 5). Analysing the performances of the logistic regression with in input the severity scores 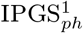 and 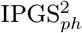 taken separately, we note that the proposed severity score models are substantially equivalent. The small differences in terms of accuracy scores within the same sample are due to the genetic algorithm procedure: when different minima, but close in the parameter space, are reached, the algorithm cannot easily escape and we accept the proposed solution as the asymptotic one. However it is worth noticing that a relevant difference remains when comparing the results obtained on the male and female data set. For females the single scores reach an accuracy of about 59%, while for the male sample we reach an excellent 76%, at the contrary with what we have seen in Fig. 6, where the logistic regression with other variables in input (such as age, IPGS) allows us to obtain similar results for the male and female sample. We can speculate that the different results in the two data samples are due to the differences in the genetic pool between male and females, since the total number of genes contributing to COVID-19 clinical variability was 4260 in males and 4360 in females, 75% of which were in common. Therefore the non-common set of genes (25%) may be determinant in giving different results. Another hypothesis is related to the fact that males are more prone to have a more severe disease respect to females, therefore we have more phenotypic data for males and more male patients analyzed (1872 males with respect to 1254 females): more specific information in this case means better training and higher performances in the testing procedure. However it is clear that the information on the organ involment is independent of the chosen severity score model and that the genetic Pygad algorithm works very well in highlighting this aspect. The proposed severity scores perform comparably within the same sample data because they are both able to convey all the relevant information from the clinical data collection, even though they are derived from different principles and they are functionally different.

### GRADING_3_

In this second part of the section we present the results related to GRADING_3_, where we have reduced the severity scores to a 3-level classification of patients into non-hospitalized (GRADING_3_ =0), hospitalized not receiving supplemental oxygen or receiving low-flow oxygen (GRADING_3_ =1) and patients with severe disease (GRADING_3_ =2). In this last case patients are considered to manifest a severe disease when they are hospitalized and receiving intensive or invasive respiratory support or are dead. In Fig. 7 are shown the confusion matrices for the male (panels (a), (b)) and female (panels (c),(d)) data set, where the best fit is presented for both sets. The results are relative to a logistic regression with multiple predictor variables used as input: age, IPGS and 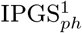 for panels (a), (c); age, IPGS and 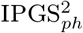 for panels (b), (d). Within the same data set, the performances of the severity scores are comparable, while, comparing between the two data sets, the accuracy experienced on the female sample is higher than the one on the male sample, irrespectively of the chosen severity score. In general, the accuracy reached in each case for GRADING_3_ is lower than the one reached for GRADING_2_ and shown in Fig. 3, due to binning limitations. If we look in detail at the confusion matrices shown in Fig. 7, it turns out that the biggest errors are done in two cases: i) when we have to predict 0 and we predict 1; ii) when we have to predict 2 and we predict 1. Probably the information that we have on the clinical framework of each patient is not optimized for distinguishing between low and severe disease, thus explaining why the GRADING_2_ was performing better, since the algorithm was not asked to distinguish between low and severe disease for GRADING_2_, but all the hospitalized patients were treated in the same way. A general comparison of the performances of the logistic regression on GRADING3 is shown in the Tables 6, 7 for the female and male data sets respectively. Analogously to the results obtained for GRADING_2_, the performances are enhanced when calculating the logistic regression on age, IPGS and 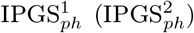, with respect to the calculation on age, IPGS only. However the enhancement is slighter with respect to what observed before and it does not hold for the case (age, IPGS and 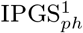) for the female data set.

**Figure 7:**
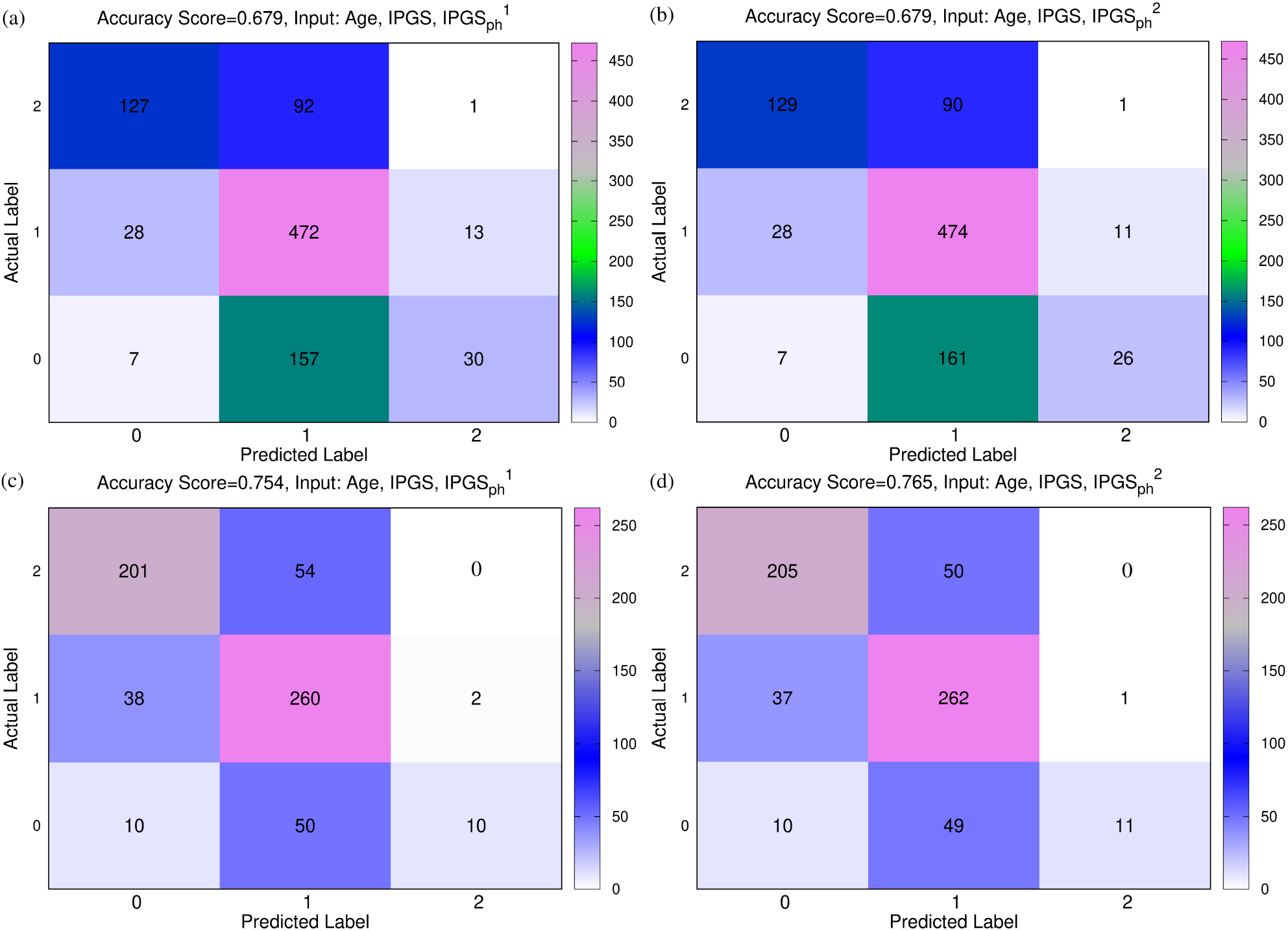
Comparison between 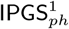 and 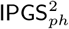. Confusion matrices obtained from a logistic regression of the single scores with age and IPGS for the male (panels (a), (b)) and female (panels (c), (d) data set. Panels (a), (c) show results for 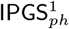; (b) (d) for 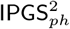

**Table 6.** Accuracy, precision, sensitivity and specificity scores resulting from a logistic regression on the female data set for GRADING_3_. The calculation of precision and sensitivity is done applying the sklearn.metrics module [34, 24] in python.

| Input Variables | Accuracy | Precision | Sensitivity | Specificity |
| --- | --- | --- | --- | --- |
| age + IPGS | 0.758 | 0.8515 | 0.603 | 0.852 |
| age + IPGS<sub>ph</sub><sup>1</sup> | 0.659 | 0.793 | 0.497 | 0.793 |
| age + IPGS<sub>ph</sub><sup>2</sup> | 0.654 | 0.789 | 0.4901 | 0.789 |
| age+ IPGS + IPGS<sub>ph</sub><sup>1</sup> | 0.754 | 0.849 | 0.633 | 0.849 |
| age+ IPGS + IPGS<sub>ph</sub><sup>2</sup> | 0.765 | 0.855 | 0.611 | 0.856 |

**Table 7.** Accuracy, precision, sensitivity and specificity scores resulting from a logistic regression on the male data set for GRADING_3_. The calculation of precision and sensitivity is done applying the sklearn.metrics module [34, 24] in python.

| Input Variables | Accuracy | Precision | Sensitivity | Specificity |
| --- | --- | --- | --- | --- |
| age + IPGS | 0.673 | 0.773 | 0.543 | 0.763 |
| age + IPGS<sup>1</sup><sub>ph</sub> | 0.585 | 0.706 | 0.406 | 0.706 |
| age + IPGS<sup>2</sup><sub>ph</sub> | 0.589 | 0.707 | 0.407 | 0.708 |
| age+ IPGS + IPGS<sup>1</sup><sub>ph</sub> | 0.679 | 0.777 | 0.551 | 0.777 |
| age+ IPGS + IPGS<sup>2</sup><sub>ph</sub> | 0.679 | 0.776 | 0.548 | 0.776 |

### GRADING_5_

We finally present the results related to GRADING_5_, where we have applied the WHO severity-grading in 6 points to classify the patients. In Fig. 8 are shown the confusion matrices for the male (panels (a), (b)) and female (panels (c),(d)) data set, where the best fit is presented for both sets. The results are relative to a logistic regression with multiple predictor variables used in input: age, IPGS and 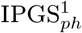 for panels (a), (c); age, IPGS and 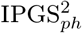 for panels (b), (d). Moreover a general comparison of the performances of the logistic regression on GRADING_5_ is shown in the Tables 8, 9 for the female and male data sets respectively.

**Figure 8:**
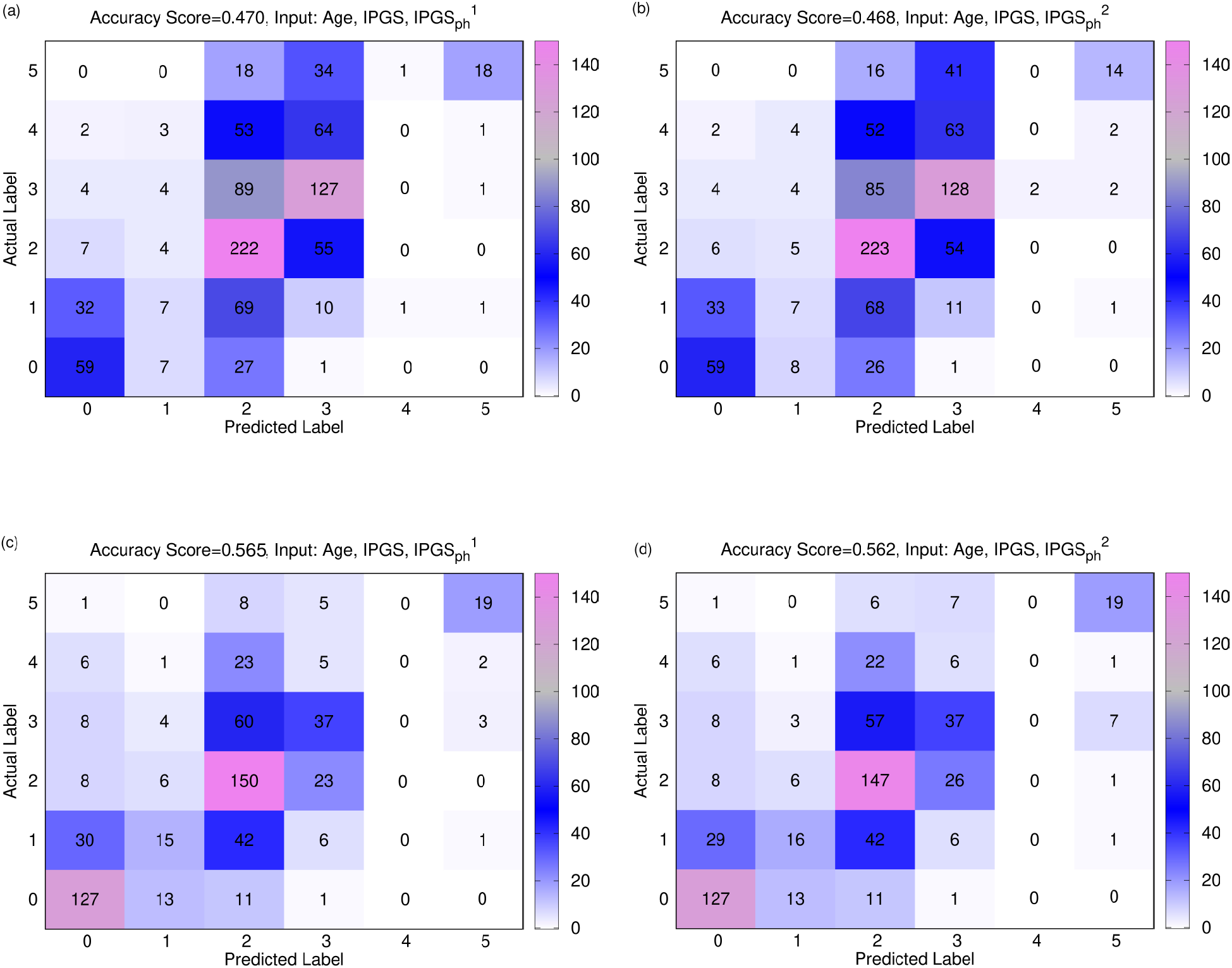
Comparison between 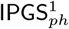 and 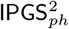. Confusion matrices obtained from a logistic regression of the single scores with age and IPGS for the male (panels (a), (b)) and female (panels (c), (d) data set. Panels (a), (c) show results for 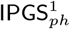; (b), (d) for 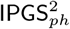

**Table 8.** Accuracy, precision and sensitivity scores resulting from a logistic regression on the female data sets for GRADING_5_. The calculation of precision and sensitivity is done applying the sklearn.metrics module [34, 24] in python.

| Input Variables | Accuracy | Precision | Sensitivity |
| --- | --- | --- | --- |
| age + IPGS | 0.560 | 0.9014 | 0.446 |
| age + IPGS<sup>1</sup><sub>ph</sub> | 0.4416 | 0.8705 | 0.2852 |
| age + IPGS<sup>2</sup><sub>ph</sub> | 0.433 | 0.869 | 0.263 |
| age+ IPGS + IPGS<sup>1</sup><sub>ph</sub> | 0.565 | 0.902 | 0.4499 |
| age+ IPGS + IPGS<sup>2</sup><sub>ph</sub> | 0.562 | 0.9021 | 0.4489 |

**Table 9.** Accuracy, precision and sensitivity scores resulting from a logistic regression on the male and female data sets for GRADING_5_. The calculation of precision and sensitivity is done applying the sklearn.metrics module [34, 24] in python.

| Input Variables | Accuracy | Precision | Sensitivity |
| --- | --- | --- | --- |
| age + IPGS | 0.460 | 0.877 | 0.3629 |
| age + IPGS<sup>1</sup><sub>ph</sub> | 0.339 | 0.8449 | 0.230 |
| age + IPGS<sup>2</sup><sub>ph</sub> | 0.332 | 0.842 | 0.2143 |
| age+ IPGS + IPGS<sup>1</sup><sub>ph</sub> | 0.470 | 0.879 | 0.379 |
| age+ IPGS + IPGS<sup>2</sup><sub>ph</sub> | 0.468 | 0.879 | 0.371 |

As already seen for GRADING_3_, within the same data set, the performances of the severity scores are comparable when predicting the GRADING_5_, while, comparing between the two data sets, the accuracy experienced on the female sample is higher than the one on the male sample, irrespectively of the chosen severity score. In general, the accuracy reached in each case for GRADING_5_ is lower than the those reached for both GRADING_2_ and GRADING_3_. If we look in detail at the confusion matrices presented in Fig. 8, the biggest errors are related to the false positive values detected for classes 2 and 3. While the algorithm seems to identify quite well the extreme classes 0 and 5, more difficulties are encountered when it comes to distinguishing between the intermediate class levels. Finally, if we compare the results of the logistic regression performed with age and IPGS as inputs with those obtained with inputs age, IPGS and 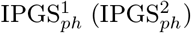, we observe a slight increase of the performances when considering two severity scores at the same time (in line with what shown for GRADING_3_ and GRADING_2_). At the contrary, a slight decrease in the performances is observed if we compare the results of the logistic regression with age and 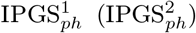 as inputs with respect to the analogous case with inputs age and IPGS (as seen also for GRADING_3_).

## 4. Conclusions

In this article we have presented two severity scores that, starting from the Integrated PolyGenic Score (IPGS) introduced in [16], integrate the phenotype of the analyzed patients in order to improve the accuracy, sensitivity and specificity performances registered by the IPGS. The performances of the proposed methods, based on a combination of clinical and genetic information, are significantly higher that the performances of methods based on genetic information alone.

However, since both our scores include information about organ involvements, that are available only in the course of the viral infection, these scores cannot be used as predictive tools in the general population, thus resulting as the main limitation of the study. Overcoming this limitation may pave the way for next generation methods able to better understand the genetic bases of complex disorders. The development of a tool able to predict, prior to viral infection, if one will be severely affected, would have a tremendous impact on the social life and world economy, improving our capability of treatment and thus reducing mortality. In this view, COVID-19 disease represents an ideal scenario for developing methods that could be used for other complex disorders since, compared to other complex disorders, in COVID-19 the environmental trigger is well known (e.g. SARS-CoV-2 infection).

## Acknowledgments

This study is part of the GEN-COVID Multicenter Study, (https://sites.google.com/dbm.unisi.it/gen-covid), the Italian multicenter study aimed at identifying the COVID-19 host genetic bases. Specimens were provided by the COVID-19 Biobank of Siena, which is part of the Genetic Biobank of Siena, member of BBMRI-IT, of Telethon Network of Genetic Biobanks (project no. GTB18001), of EuroBioBank, and of RD-Connect. All authors of this paper are members of European Reference Network on rare respiratory diseases ERN-LUNG. We thank the CINECA consortium for providing computational resources and the Network for Italian Genomes (NIG; http://www.nig.cineca.it) for its support. We thank private donors for the support provided to AR (Department of Medical Biotechnologies, University of Siena) for the COVID-19 host genetics research project (D.L n.18 of March 17, 2020). We also thank the COVID-19 Host Genetics Initiative (https://www.covid19hg.org/), MIUR project “Dipartimenti di Eccellenza 2018-2020” to the Department of Medical Biotechnologies University of Siena, Italy, and “Bando Ricerca COVID-19 Toscana” project to Azienda Ospedaliero-Universitaria Senese. We thank Intesa San Paolo for the 2020 charity fund dedicated to the project N B/2020/0119 “Identificazione delle basi genetiche determinanti la variabilità clinica della risposta a COVID-19 nella popolazione italiana”. The Italian Ministry of University and Research for funding within the “Bando FISR 2020” in COVID-19 fo the project “Editing dell’RNA contro il Sars-CoV-2: hackerare il virus per identificare bersagli molecolari e attenuare l’infezione - HACKTHECOV” and the Istituto Buddista Italiano Soka Gakkai for funding the project “PAT-COVID: Host genetics and pathogenetic mechanisms of COVID-19” (ID n. 2020-2016_RIC_3). We thank EU project H2020-SC1-FA-DTS-2018-2020, entitled “International consortium for integrative genomics prediction (INTERVENE)” - Grant Agreement No. 101016775. Generous support was also received from private donations by Mrs. Maurizio Traglio, Enzo Cattaneo and Alberto Borella.

## GEN-COVID Multicenter study (https://sites.google.com/dbm.unisi.it/gen-covid)

Francesca Mari^1,2,3^, Sergio Daga^1,2^, Ilaria Meloni^1,2^, Mirella Bruttini^1,2,3^, Susanna Croci^1,2^, Mirjam Lista^1,2^, Debora Maffeo^1,2^, Elena Pasquinelli^1,2^, Giulia Brunelli^1,2^, Kristina Zguro^1,2^, Viola Bianca Serio^1,2^, Enrica Antolini^1,2^Simona Letizia Basso^1,2^, Samantha Minetto^1,2^, Giulia Rollo^1,2^, Martina Rozza^1,2^, Angela Rina^1,2^, Rossella Tita^3^, Maria Antonietta Mencarelli^3^, Caterina Lo Rizzo^3^, Anna Maria Pinto^3^, Francesca Ariani^1,2,3^, Francesca Montagnani^2,4^, Mario Tumbarello^2,4^, Ilaria Rancan^2,4^, Massimiliano Fabbiani^4^, Elena Bargagli^5^, Laura Bergantini^5^, Miriana d’Alessandro^5^, Paolo Cameli^5^, David Bennett^5^, Federico Anedda^6^, Simona Marcantonio^6^, Sabino Scolletta^6^, Federico Franchi^6^, Maria Antonietta Mazzei^7^, Susanna Guerrini^7^, Edoardo Conticini^8^, Luca Cantarini^8^, Bruno Frediani^8^, Danilo Tacconi^9^, Chiara Spertilli Raffaelli^9^, Arianna Emiliozzi^9^, Marco Feri^10^, Alice Donati^10^, Raffaele Scala^11^, Luca Guidelli^11^, Genni Spargi^12^, Marta Corridi^12^, Cesira Nencioni^13^, Leonardo Croci^13^, Gian Piero Caldarelli^14^, Davide Romani^15^, Paolo Piacentini^15^, Maria Bandini^15^, Elena Desanctis^15^, Silvia Cappelli^15^, Anna Canaccini^16^, Agnese Verzuri^16^, Valentina Anemoli^16^, Manola Pisani^16^, Agostino Ognibene Maria Lorubbio^17^, Alessandro Pancrazzi^17^, Massimo Vaghi^18^, Antonella D’Arminio Monforte^19^, Federica Gaia Miraglia^19^, Mario U. Mondelli^20,21^, Stefania Mantovani^20^, Raffaele Bruno^20,22^, Marco Vecchia^20^, Marcello Maffezzoni^22^, Enrico Martinelli^23^, Massimo Girardis^24^, Stefano Busani^24^, Sophie Venturelli^24^, Andrea Cossarizza^25^, Andrea Antinori^26^, Alessandra Vergori^26^, Stefano Rusconi^27,28^, Matteo Siano^28^, Arianna Gabrieli^28^, Agostino Riva^27,28^, Daniela Francisci^29^, Elisabetta Schiaroli^29^, Carlo Pallotto^29^, Saverio Giuseppe Parisi^30^, Monica Basso^30^, Sandro Panese^31^, Stefano Baratti^31^, Pier Giorgio Scotton^32^, Francesca Andretta^32^, Mario Giobbia^32^, Renzo Scaggiante^33^, Francesca Gatti^33^, Francesco Castelli^34^, Eugenia Quiros-Roldan^34^, Melania Degli Antoni^34^, Isabella Zanella^35,36^, Matteo della Monica^37^, Carmelo Piscopo^37^, Mario Capasso^38,39^, Roberta Russo^38,39^, Immacolata Andolfo^38,39^, Achille Iolascon^38,39^, Giuseppe Fiorentino^40^, Massimo Carella^41^, Marco Castori^41^, Giuseppe Merla^38,42^, Gabriella Maria Squeo^42^, Filippo Aucella^43^, Pamela Raggi^44^, Rita Perna^44^, Matteo Bassetti^45,46^, Antonio Di Biagio^45,46^, Maurizio Sanguinetti^47,48^, Luca Masucci^47,48^, Alessandra Guarnaccia^47^, Serafina Valente^49^, Alex Di Florio^49^, Marco Mandalà^50^, Alessia Giorli^50^, Lorenzo Salerni^50^, Patrizia Zucchi^51^, Pierpaolo Parravicini^51^, Elisabetta Menatti^52^, Tullio Trotta^53^, Ferdinando Giannattasio^53^, Gabriella Coiro^53^, Fabio Lena^54^, Gianluca Lacerenza^54^, Cristina Mussini^55^, Luisa Tavecchia^56^, Lia Crotti^57,58,59,60,61^, Gianfranco Parati^57,58^, Roberto Menè^57,58^, Maurizio Sanarico^62^, Marco Gori^63,64^, Francesco Raimondi^65^, Alessandra Stella^65^, Filippo Biscarini^66^, Tiziana Bachetti^67^, Maria Teresa La Rovere^68^, Maurizio Bussotti^69^, Serena Ludovisi^70^, Katia Capitani^71^, Simona Dei^72^, Sabrina Ravaglia^73^, Annarita Giliberti^74^, Giulia Gori^74^, Rosangela Artuso^74^, Elena Andreucci^74^, Angelica Pagliazzi^74^, Erika Fiorentini^74^, Antonio Perrella^75^, Francesco Bianchi^2,75^, Paola Bergomi^76^, Emanuele Catena^76^, Riccardo Colombo^76^, Sauro Luchi^77^, Giovanna Morelli^77^, Paola Petrocelli^77^, Sarah Iacopini^77^, Sara Modica^77^, Silvia Baroni^78^, Giulia Micheli^79^, Marco Falcone^80^, Donato Urso^80^, Giusy Tiseo^80^, Tommaso Matucci^80^, Davide Grassi8^1^, Claudio Ferri^81^, Franco Marinangeli^82^, Francesco Brancati^83,84^, Antonella Vincenti^85^, Valentina Borgo^85^, Stefania Lombardi^85^, Mirco Lenzi^85^, Massimo Antonio Di Pietro^86^, Francesca Vichi^86^, Benedetta Romanin^86^, Letizia Attala^86^, Cecilia Costa^86^, Andrea Gabbuti^86^, Alessio Bellucci^86^, Marta Colaneri^22^, Patrizia Casprini^87^, Cristoforo Pomara^88^, Massimiliano Esposito^88^, Roberto Leoncini^89^, Michele Cirianni^89^, Lucrezia Galasso^89^, Marco Antonio Bellini^90^, Chiara Gabbi^91^, Nicola Picchiotti^63^

1) Medical Genetics, University of Siena, Siena, 53100, Italy

2) Med Biotech Hub and Competence Center, Department of Medical Biotechnologies, University of Siena, Siena, 53100, Italy

3) Genetica Medica, Azienda Ospedaliero-Universitaria Senese, Siena, 53100, Italy

4) Department of Medical Sciences, Infectious and

Tropical Diseases Unit, Azienda Ospedaliera Universitaria Senese, Siena, 53100, Italy

5) Unit of Respiratory Diseases and Lung Transplantation, Department of Internal and Specialist Medicine, University of Siena, Siena, 53100, Italy

6) Department of Emergency and Urgency, Medicine, Surgery and Neurosciences, Unit of Intensive Care Medicine, Siena University Hospital, Siena, 53100, Italy

7) Department of Medical, Surgical and Neuro Sciences and Radiological Sciences, Unit of Diagnostic Imaging, University of Siena, 53100, Italy

8) Rheumatology Unit, Department of Medicine, Surgery and Neurosciences, University of Siena, Policlinico Le Scotte, 53100, Italy

9) Department of Specialized and Internal Medicine, Infectious Diseases Unit, San Donato Hospital Arezzo, Italy

10) Department of Emergency, Anesthesia Unit, San Donato Hospital, Arezzo, Italy

11) Department of Specialized and Internal Medicine, Pneumology Unit and UTIP, San Donato Hospital, Arezzo, Italy

12) Department of Emergency, Anesthesia Unit, Misericordia Hospital, Grosseto, Italy

13) Department of Specialized and Internal Medicine, Infectious Diseases Unit, Misericordia Hospital, Grosseto, Italy

14) Clinical Chemical Analysis Laboratory, Misericordia Hospital, Grosseto, Italy

15) Dipartimento di Prevenzione, Azienda USL Toscana Sud Est, Italy

16) Dipartimento Tecnico-Scientifico Territoriale, Azienda USL Toscana Sud Est, Italy

17) UOC Laboratorio Analisi Chimico Cliniche, Arezzo, Italy

18) Chirurgia Vascolare, Ospedale Maggiore di Crema, Italy

19) Department of Health Sciences, Clinic of Infectious Diseases, ASST Santi Paolo e Carlo, University of Milan, Italy

20) Division of Clinical Immunology - Infectious Diseases, Department of Medicine, Fondazione IRCCS Policlinico San Matteo, Pavia, Italy

21) Department of Internal Medicine and Therapeutics, University of Pavia, Italy

22) University of Pavia, Pavia, Italy

23) Department of Respiratory Diseases, Azienda Ospedaliera di Cremona, Cremona, Italy

24) Department of Anesthesia and Intensive Care, University of Modena and Reggio Emilia, Modena, Italy

25) Department of Medical and Surgical Sciences for Children and Adults, University of Modena and Reggio Emilia, Modena, Italy

26) HIV/AIDS Department, National Institute for Infectious Diseases, IRCCS, Lazzaro Spallanzani, Rome, Italy

27) III Infectious Diseases Unit, ASST-FBF-Sacco, Milan, Italy

28) Department of Biomedical and Clinical Sciences Luigi Sacco, University of Milan, Milan, Italy

29) Infectious Diseases Clinic, “Santa Maria della Misericordia” Hospital, University of Perugia, Perugia, Italy

30) Department of Molecular Medicine, University of Padova, Italy

31) Clinical Infectious Diseases, Mestre Hospital, Venezia, Italy

32) Department of Infectious Diseases, Treviso Hospital, Local Health Unit 2 Marca Trevigiana, Treviso, Italy

33) Infectious Diseases Clinic, ULSS1, Belluno, Italy

34) Department of Infectious and Tropical Diseases, University of Brescia and ASST Spedali Civili Hospital, Brescia, Italy

35) Department of Molecular and Translational Medicine, University of Brescia, Italy

36) Clinical Chemistry Laboratory, Cytogenetics and Molecular Genetics Section, Diagnostic Department, ASST Spedali Civili di Brescia, Italy

37) Medical Genetics and Laboratory of Medical Genetics Unit, A.O.R.N. “Antonio Cardarelli”, Naples, Italy

38) Department of Molecular Medicine and Medical Biotechnology, University of Naples Federico II, Naples, Italy

39) CEINGE Biotecnologie Avanzate, Naples, Italy

40) Unit of Respiratory Physiopathology, AORN dei Colli, Monaldi Hospital, Naples, Italy

41) Division of Medical Genetics, Fondazione IRCCS Casa Sollievo della Sofferenza Hospital, San Giovanni Rotondo, Italy

42) Laboratory of Regulatory and Functional Genomics, Fondazione IRCCS Casa Sollievo della Sof-ferenza

43) Department of Medical Sciences, Fondazione IRCCS Casa Sollievo della Sofferenza Hospital, San Giovanni Rotondo, Italy

44) Clinical Trial Office, Fondazione IRCCS Casa Sollievo della Sofferenza Hospital, San Giovanni Rotondo, Italy

45) Department of Health Sciences, University of Genova, Genova, Italy

46) Infectious Diseases Clinic, Policlinico San Martino Hospital, IRCCS for Cancer Research Genova, Italy

47) Microbiology, Fondazione Policlinico Universitario Agostino Gemelli IRCCS, Catholic University of Medicine, Rome, Italy

48) Department of Laboratory Sciences and Infectious Diseases, Fondazione Policlinico Universitario A. Gemelli IRCCS, Rome, Italy

49) Department of Cardiovascular Diseases, University of Siena, Siena, Italy

50) Otolaryngology Unit, University of Siena, Italy

51) Department of Internal Medicine, ASST Valtellina e Alto Lario, Sondrio, Italy

52) Study Coordinator Oncologia Medica e Ufficio Flussi Sondrio, Italy

53) First Aid Department, Luigi Curto Hospital, Polla, Salerno, Italy

54) Department of Pharmaceutical Medicine, Misericordia Hospital, Grosseto, Italy

55) Infectious Diseases Clinics, University of Modena and Reggio Emilia

56) U.O.C. Medicina, ASST Nord Milano, Ospedale Bassini, Cinisello Balsamo (MI), Italy

57) IRCCS, Istituto Auxologico Italiano, Dipartimento di Cardiologia, Ospedale San Luca, Milano, Italy

58) Department of Medicine and Surgery, University of Milano-Bicocca, Milan, Italy

59) Istituto Auxologico Italiano, IRCCS, Center for Cardiac Arrhythmias of Genetic Origin, Milan, Italy

60) Istituto Auxologico Italiano, IRCCS, Laboratory of Cardiovascular Genetics, Milan, Italy

61) Member of the European Reference Network for Rare, Low Prevalence and Complex Diseases of the Heart-ERN GUARD-Heart

62) Independent Data Scientist, Milan, Italy

63) University of Siena, DIISM-SAILAB, Siena, Italy

64) Maasai, I3S CNRS, Université Côte d’Azur, France

65) Laboratorio di Biologia Bio@SNS, Scuola Normale Superiore, Pisa, Italy

66) CNR-Consiglio Nazionale delle Ricerche, Istituto di Biologia e Biotecnologia Agraria (IBBA), Milano, Italy

67) Direzione Scientifica, Istituti Clinici Scientifici Maugeri IRCCS, Pavia, Italy

68) Istituti Clinici Scientifici Maugeri IRCCS, Department of Cardiology, Institute of Montescano, Pavia, Italy

69) Istituti Clinici Scientifici Maugeri IRCCS, Department of Cardiology, Institute of Milan, Italy

70) Fondazione IRCCS Ca’ Granda Ospedale Maggiore Policlinico, Milan, Italy

71) Core Research Laboratory, ISPRO, Florence, Italy

72) Health Management, Azienda USL Toscana Sud Est, Tuscany, Italy

73) IRCCS C. Mondino Foundation, Pavia, Italy

74) Medical Genetics Unit, Meyer Children’s University Hospital

75) Department of Medicine, Pneumology Unit, Misericordia Hospital, Grosseto, Italy

76) Department of Anesthesia and Intensive Care Unit, ASST Fatebenefratelli Sacco, Luigi Sacco Hospital, Polo Universitario, University of Milan, Milan

77) Infectious Disease Unit, Hospital of Lucca, Italy

78) Department of Diagnostic and Laboratory Medicine, Institute of Biochemistry and Clinical Biochemistry, Fondazione Policlinico Universitario A. Gemelli IRCCS, Catholic University of the Sacred Heart, Rome, Italy

79) Clinic of Infectious Diseases, Catholic University of the Sacred Heart, Rome, Italy

80) Department of Clinical and Experimental Medicine, Infectious Diseases Unit, University of Pisa, Pisa, Italy

81) Department of Clinical Medicine, Public Health, Life and Environment Sciences, University of L’Aquila, Italy

82) Anesthesiology and Intensive Care, University of L’Aquila, L’Aquila, Italy

83) Department of Life, Health and Environmental Sciences, University of L’Aquila, 67100, L’Aquila, Italy

84) Human Functional Genomics Laboratory, IRCCS San Raffaele Roma, 00167, Rome, Italy

85) Infectious Disease Unit, Hospital of Massa, Italy

86) Infectious Diseases Unit, Santa Maria Annunziata Hospital, USL Centro, Florence, Italy

87) Laboratory of Clinical Pathology and Immunoallergy, Florence-Prato, Italy

88) Department of Medical, Surgical and Advanced Technologies “G.F. Ingrassia”, University of Catania, Catania, Italy

89) Laboratorio Patologia Clinica, Azienda Ospedaliero-Universitaria Senese

90) Ambulatory Chronic Polipathology of Siena, Department of Medicine, Surgery and Neurosciences, University of Siena, Siena, Italy

91) Department of Biosciences and Nutrition, Karolinska Institutet, Stockholm, Sweden

